# Environmental factors and host sex influence the skin microbiome structure of Hong Kong newt (*Paramesotriton hongkongensis*) in a coldspot of chytridiomycosis in subtropical East Asia

**DOI:** 10.1101/2024.02.19.581002

**Authors:** Bowen Wan, Guoling Chen, Emily Shui Kei Poon, Hon Shing Fung, Anthony Lau, Simon Yung Wa Sin

## Abstract

Chytridiomycosis, an infectious skin disease caused by the chytrid fungi, *Batrachochytrium dendrobatidis* and *B. salamandrivorans*, has been threatening global amphibian biodiversity. On the skin of chytrid-resistant amphibians, some antifungal bacteria likely defend them against chytridiomycosis, reducing the mortality of resistant amphibians. The Hong Kong newt (*Paramesotriton hongkongensis*) inhabits East Asia which is a suspected origin of chytrids. Asymptomatic infection in the newts suggests their long-term coexistence with chytrids. Thus, the skin microbiomes of this resistant species warrant investigation, along with other factors that can affect the microbiome. Among the 149 newts sampled in their natural habitats in Hong Kong, putative antifungal bacteria were found on all newts. There were 314 amplicon sequence variants distributed over 25 genera of putative antifungal bacteria, abundant ones included *Acinetobacter*, *Flavobacterium*, and *Novosphingobium* spp. The skin microbiome compositions were strongly influenced by the inter-site geographical distances. Despite inter-site differences, we identified some core skin microbes across sites, which could be vital to *P. hongkongensis*. The dominant cores included family Comamonadaceae, family Chitinophagaceae, and class Betaproteobacteria. Moreover, habitat elevation and host sex also exhibited significant effects on skin microbiome compositions. The antifungal bacteria found on these newts offer an important resource for conservation against chytridiomycosis, such as probiotic treatments for susceptible species.

## Introduction

Microbiomes, communities of microorganisms, have been found to reside on or inside animals (Hoffmann, 2017). Symbionts in the microbiome can enhance the innate immunity of the host by educating the immune system (Naik et al., 2015, Chen and Tsao, 2013) and help the host combat infections through competition for food and space with pathogenic microbes or through the production of antimicrobial compounds against invasive pathogens (Grice and Segre, 2011, Chen and Tsao, 2013). It has been shown in germ-free laboratory mice that the lack of skin microbiomes led to compromised immunity involving aberrant epidermal T cell populations, which could be rescued by the colonization of commensal skin bacteria, e.g., *Staphylococcus epidermidis* (Chen and Tsao, 2013, Naik et al., 2015). In humans, fecal microbiota transplant is a treatment for patients with recurrent *Clostridioides difficile* infections using a fecal preparation from a healthy stool donor to repopulate the patient guts with balanced microbiota that compete with *C. difficile* for space and resources (Cammarota et al., 2014). These cases highlight the crucial role of resident body microbes in maintaining the host’s health, indicating a coevolutionary relationship between these organisms and their host (Colston and Jackson, 2016, Naik et al., 2015). This hypothesis of coevolution is supported by the observation of phylosymbiosis, in which there is a significant cophylogenetic signal between the microbiomes and their corresponding hosts (Brooks et al., 2016, Ross et al., 2019, Colston and Jackson, 2016).

Among vertebrates, amphibians are the most endangered group, with 41% of all species threatened with extinction (IUCN, 2021). The causes of amphibian population decline are numerous and often complex. Chytridiomycosis is an infection that has caused the most significant biodiversity loss among diseases, contributing to the decline of at least 501 amphibian species, of which 90 are extinct, and 124 have declined by over 90% in abundance (Fisher and Garner, 2020, Scheele et al., 2019). For example, in a study conducted in Panama, 30 amphibian species were found to be lost after the epidemic outbreak of chytridiomycosis, including three caudate species *Bolitoglossa colonnea*, *B. schizodactyla*, and *Oedipina parvipes* (Crawford et al., 2010). Before the identification of *Batrachochytrium salamandrivorans* (*Bsal*) in 2013, chytridiomycosis was believed to be caused solely by the pathogenic chytrid fungus, *Batrachochytrium dendrobatidis* (*Bd*), leaving some dramatic amphibian population declines unexplained, including the local extinction of the fire salamander (*Salamandra salamandra*) in the Netherlands (Spitzen-van der Sluijs et al., 2013, Martel et al., 2013). It has been subsequently discovered that *Bsal* is a salamander-and-newt-specific pathogen, unlike *Bd* which is capable of infecting a wider range of amphibian species, including frogs, salamanders, and caecilians (Martel et al., 2013, Martel et al., 2014, Fisher and Garner, 2020).

Both *Bd* and *Bsal* infect the amphibian skin, which is the organ responsible for gaseous exchange and osmoregulation with specific adaptations to aquatic and terrestrial habitats (Fisher and Garner, 2020, Varga et al., 2019). Being mucosal, water-permeable, and in direct contact with the microbially rich and diverse environment, the amphibian skin is susceptible to pathogen invasion (Varga et al., 2019). As in other vertebrates (Martín-Platero et al., 2006, Martín-Vivaldi et al., 2010, Fredricks, 2001, Lai et al., 2010, Naik et al., 2015, Pérez-Sánchez et al., 2011), microbiome is one of the vital innate immune defenses of the amphibian skin (Varga et al., 2019). Potential coevolution might have shaped a skin microbiome that confers higher resistance to infection in amphibians (Foster et al., 2017, Bates et al., 2018, Colston and Jackson, 2016). For conservation purposes, research has been focused on the amphibian skin microbiome with an emphasis on its implications for understanding chytrid pathogenesis and treatment. Previous studies have demonstrated interactions existed between skin microbiome and *Bd* infection. For example, in the Sierra Nevada yellow-legged frog (*Rana sierrae*), *Bd* was shown to disturb the skin bacterial community both in the wild populations and controlled experiments (Jani and Briggs, 2014). In the red-backed salamander (*Plethodon cinereus*), *Bd* load was found to change with skin associated microbes under temperature variation (Muletz-Wolz et al., 2019). *Bd* infection history also had a significant effect on the skin microbiome of the common midwife toad (*Alytes obstetricans*), reducing the bacterial diversity in epizootic populations (Bates et al., 2018). On the other hand, the inoculation of anti-*Bd* bacterial isolates identified from resistant species is able to reduce the morbidity, mortality as well as severity of symptoms in the mountain yellow-legged frog (*Rana muscosa*) (Harris et al., 2009a), the western toad (*Anaxyrus boreas*) (Kueneman et al., 2016), and *P. cinereus* (Harris et al., 2009b) when they are infected with *Bd*.

Contrary to the intensive research associated with *Bd*, far less is known about the emerging fungal pathogen *Bsal*, including its genetic and phenotypic variations, virulence, and threats to global salamander and newt populations (Fisher and Garner, 2020). Genetic evidence suggests that *Bd* originates in East Asia (Fisher and Garner, 2020, O’hanlon et al., 2018). It is noteworthy that *Bsal* has also only been found in East and Southeast Asia, apart from the epidemic in Europe (Yuan et al., 2018, Laking et al., 2017, Martel et al., 2014). Referred as a coldspot of chytridiomycosis, Asia has a low prevalence of chytrids and lacked chytridiomycosis outbreaks, which hints at the coevolution of chytrids and Asian amphibians (Swei et al., 2011, Laking et al., 2017, Yuan et al., 2018). However, amphibian skin microbiome studies in Asia are scarce relative to Europe and North America. It has been found in European newt species, including the smooth newt (*Lissotriton vulgaris*), the great-crested newt (*Triturus cristatus*), and *S. salamandra*, that *Bsal* infection was associated with altered skin microbial composition (Bates et al., 2019, Bletz et al., 2018). *Bsal*-inhibiting bacteria have been identified from *S. salamandra* skin microbiome, which could slow the progression of disease chytridiomycosis when increased in abundance (Bletz et al., 2018).

Besides chytrid fungal infection, the amphibian microbial community is also affected by various host and environmental factors. Host species identity was proposed as the strongest predictor of the composition of a skin microbial community (Kueneman et al., 2014). This observation is supported by other studies, such as the one that documented different Panamanian frog species harboring distinct skin bacterial communities (Belden et al., 2015). Moreover, the effect of life history traits was observed in *A. boreas* (Prest et al., 2018) and the Cascades frog (*Rana cascadae*) (Kueneman et al., 2017, Kueneman et al., 2014) as their skin microbial compositions experienced major changes from tadpole to adult stage. While the effect of host sex on the skin microbiome is not well documented in amphibian studies, its influence was nonetheless found in a plethodontid salamander, the yellow-eyed ensatina (*Ensatina eschscholtzii xanthoptica*) (Prado-Irwin et al., 2017). However, in other plethodontid species, skin microbiome structures are more influenced by geographical parameters such as elevation and habitat type rather than phylogenetic relatedness or life stages (Prado-Irwin et al., 2017, Bird et al., 2018, Muletz Wolz et al., 2018), as environmental factors often co-vary along the elevation gradient of a habitat (Muletz Wolz et al., 2018, Hughey et al., 2017). Similar findings were reported in the cane toad (*Rhinella marina*), in which geographical location dictated the structure of the skin microbial community (Abarca et al., 2018). In addition, co-evolution with predators might also drive the recruitment and proliferation of toxin-producing bacteria in the skin microbiomes of amphibians (Vaelli et al., 2020). A recent study reported that the bacteria from the genera *Aeromonas*, *Pseudomonas*, *Shewanella*, and *Sphingopyxis* on the rough-skinned newt (*Taricha granulosa*) produce a deadly neurotoxin, tetrodotoxin (TTX), which could defend the host against predation (Vaelli et al., 2020). Nevertheless, the relative roles of different factors in affecting amphibian skin microbiomes are still unclear and are likely context-dependent.

The Hong Kong newt (*Paramesotriton hongkongensis*) (Fig. 1a) is a subtropical salamander listed as Near Threatened by the International Union for Conservation of Nature (IUCN) with a highly restricted distribution in southern China (IUCN, 2021, Fu et al., 2013, Lau et al., 2017, Lau, 2017). A 2018 screening survey of 36 salamander species in China found that *P. hongkongensis* had the highest incidence of *Bsal* presence on the skin at >50%, with all *Bsal*-positive individuals found in a single population in Wutongshan near Hong Kong (Yuan et al., 2018). Given the low number of skin microbiome studies on amphibians, specifically salamanders, in Asia in relation to the origins of *Bd* and *Bsal*, the knowledge generated from this study can offer important insights into the host-pathogen-microbiome coevolution in the region. Furthermore, the anti-*Bd* bacterial communities found on the salamander skin in this chytrid-coldspot region can be a useful resource for amphibian conservation, such as providing targets for developing probiotic treatments against chytridiomycosis. The high *Bsal* prevalence and absence of host symptoms reported in the previous survey make *P. hongkongensis* a suitable system for these purposes (Yuan et al., 2018). Here we aimed to: 1) characterize the skin microbiome compositions and the core microbiome of *P. hongkongensis* populations, 2) investigate whether *P. hongkongensis* harbors putative antifungal bacteria, and 3) determine the host and environmental factors associated with the skin microbial community structure.

**Figure 1.**
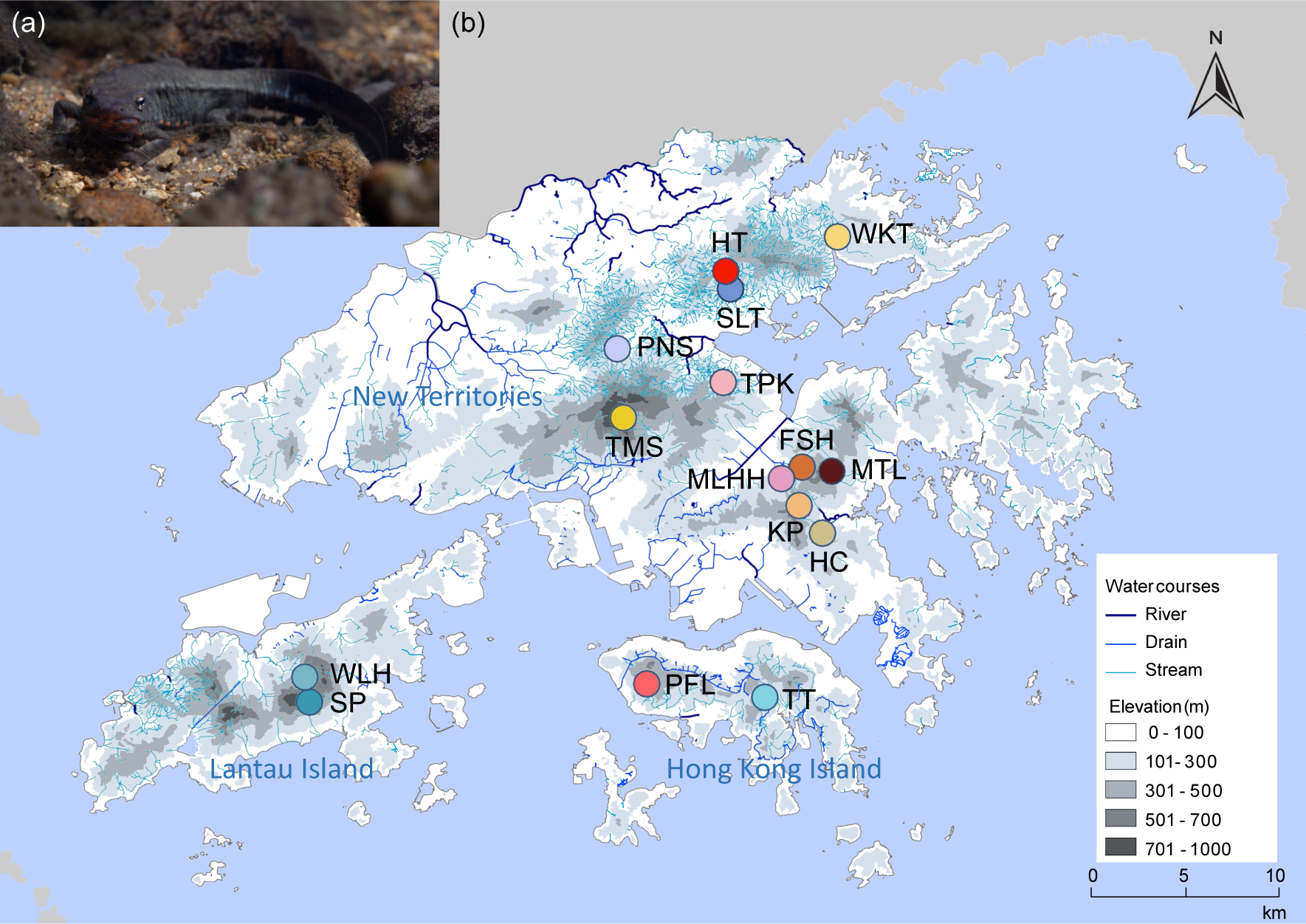
(a) *P. hongkongensis* under water. Photo credit: Hon Shing Fung. (b) Map of Hong Kong showing New Territories, Lantau Island, and Hong Kong Island, with sampling sites indicated by colored spots. Elevation is represented by shades of gray. Stream system is shown by blue lines. Abbreviations of sampling sites: FSH, Fa Sam Hang; HC, Ho Chung; HT, Hok Tau; KP, Kowloon Peak; MLHH, Ma Lai Hau Hang; MTL, Mui Tsz Lam; PNS, Pak Ngau Shek; PFL, Pok Fu Lam; SLT, Sha Lo Tung; SP, Sunset Peak; TMS, Tai Mo Shan; TPK, Tai Po Kau; TT, Tai Tam; WLH, Wong Lung Hang; and WKT, Wu Kau Tang.

## Materials and Methods

### Field sampling

We collected *P. hongkongensis* skin swabs (n=149) and water samples (n=12) from 15 locations in Hong Kong (Fig. 1b) between October 2019 and February 2020 (Supplementary Table S1), during their breeding season as the newts gathered in streams to reproduce. We captured each newt with gloved hands and kept it in an individual plastic bag. A sterile swab (FLOQSwabs, Copan, Italy) was used to stroke each individual in a single direction 30 times each on the ventral and dorsal sides and five times on each leg. We recorded the snout-to-vent length (SVL), total length (TL), weight, and sex, and calculated the body condition index (BCI, weight/SVL) for each individual. One milliliter of stream water was taken from each of the 12 locations. Samples were kept on ice immediately in the field and transferred back to the laboratory within 3 hours, then stored at −80°C until DNA extraction. This study was approved by the University animal ethics committee (EC052/2021) and the Department of Health [(19-103) in DH/HT&A/8/2/6 Pt. 1)].

### DNA extraction, DNA library preparation, and sequencing

We extracted DNA from skin swabs and water samples using the E.Z.N.A. Tissue DNA Kit (Omega Bio-tek, USA) following the manufacturer’s protocol under aseptic conditions. Four negative controls were included during DNA extraction. We also prepared mock communities (MCs) by pooling equimolar genomic DNA of multiple bacteria (Supplementary Table S2).

For high-throughput sequencing of the V4 region of the 16S rRNA gene to characterize the skin microbiome of *P. hongkongensis*, all samples, MCs, and negative controls were subjected to a two-step PCR (Cruaud et al., 2017, Huang et al., 2022) for library preparation. The first-step PCR was performed in a 30-μL reaction containing 6 μL 5X GoTaq Flexi Buffer (Promega, USA), 0.6 μL 10 mM dNTP Mix, 2.4 μL 25 mM MgCl2, 0.15 μL GoTaq G2 Flexi DNA Polymerase (Promega, USA), 1 μL DNA, 1.5 μL each of 10 μM forward and reverse primers, 3 μL 10% dimethyl sulfoxide (DMSO) (Sigma, USA), 0.15 μL 20 mg mL^-1^ bovine serum albumin (BSA) (New England Biolabs, USA), and 13.7 μL ultrapure water. The forward and reverse primers comprise 515F (Parada) (5’-GTGYCAGCMGCCGCGGTAA-3’) (Parada et al., 2016) and 806R (Apprill) (5’-GGACTACNVGGGTWTCTAAT-3’) (Apprill et al., 2015), respectively, a heterogeneity spacer and a linker. Thermocycling conditions were 95℃ for 2 min, followed by 25 cycles of 95℃ for 30 s, 50℃ for 30 s, 72℃ for 30 s, and a final extension of 72℃ for 5 min.

Products from the first-step PCR were purified and used as the template in the second-step PCR, in a 45-μL reaction consisting of 9 μL 5X GoTaq Flexi Buffer (Promega, USA), 0.9 μL 10 mM dNTP Mix, 5.4 μL 25 mM MgCl2, 0.225 μL GoTaq G2 Flexi DNA Polymerase (Promega, USA), 18.5 μL purified DNA, 0.9 μL each of 10 μM forward and 10μM reverse primers, 4.5 μL 10% DMSO, and 4.675 μL ultrapure water. The forward and reverse primers consist of the linker, an i5 or i7 index, and the adapter sequence (Cruaud et al., 2017). Each sample had a unique combination of the i5 and i7 indices. Thermocycling conditions were 95℃ for 2 min, followed by 10 cycles of 95℃ for 30 s, 55℃ for 30 s, 72℃ for 45 s, and a final extension of 72℃ for 5 min. The PCR products from both PCR steps were purified with E.Z.N.A. MicroElute Gel Extraction Kit (Omega Bio-tek, USA). The purified products from the second PCR step were quantified with the dsDNA HS Assay Kit and the Qubit 4 fluorometer (Invitrogen, USA), and pooled in an equimolar ratio to prepare a library multiplex. DNA was sequenced using NovaSeq (PE 250 bp; Illumina) by Novogene Corporation.

### Sequence data processing

De-multiplexed 16S paired-end sequence reads were merged, trimmed to remove adapters, and quality-filtered. We merged the paired-end reads with -fastq_mergepairs in USEARCH v11.0.667 (Edgar, 2010) and trimmed the primer sequences using CUTADAPT v2.4 (max_error_rate = 0.15) (Martin, 2011). Quality assessment was done with FastQC v0.11.8 (Andrews, 2010) and -fastq_eestats2 in VSEARCH v2.15.1 (Rognes et al., 2016), and only the reads with lengths between 240 and 253 bp with expected errors per read less than one were retained with the VSEARCH -fastq_filter. Pre-processed reads were then de-replicated with VSEARCH -derep_fulllength. The chimeras and singletons were removed with USEARCH -unoise3, which represented less than 0.0001% of total reads (Edgar and Flyvbjerg, 2015, Edgar, 2016b), and the remaining sequences were clustered at a 99% similarity level with VSEARCH - usearch_global to generate the amplicon sequence variants (ASVs). Taxonomy was assigned to each ASV to the lowest identifiable taxonomic level by the SINTAX algorithm of USEARCH (Edgar, 2016a) with bootstrap support higher than 80%, using the EzBioCloud 16S database as reference (assess in July 2021) (Yoon et al., 2017). The same approach was applied to identify putative antifungal ASVs using the anti-*Bd* bacteria database as reference (inhibitory ASVs only; assessed in November 2021) (Woodhams et al., 2015) with a bootstrap cutoff of 99%, and only ASVs able to be identified at the genus or species level were considered to match the sequences from putative anti-*Bd* taxa. ASVs with unidentified taxonomy or determined as contaminants according to negative controls were removed (Supplementary Table S3). Those ASVs represented by less than 0.018% [a threshold determined by MCs, as in (Huang et al., 2021); Supplementary Table S3] of total reads in each sample were considered false positives and filtered out.

### Data analysis

All statistical analyses were done in R version 4.1.0 (R Development Core Team, 2021). The microbiome composition was presented using the relative read abundance (RRA) and weighted percentage of occurrence (wPOO) of each bacterial taxon in an individual, and frequency of occurrence (FOO) of each taxon in all swab samples. Taxa with an FOO of over 90% were identified as the core microbiome. Microbiome diversity was calculated by the effective number of ASVs (hill number) for each sample (alpha diversity) and sampling site (gamma diversity) using the hilldiv package (Alberdi and Gilbert, 2019). The hill number was dependent on the order of the diversity *q* (Hill, 1973). As *q* increased, more effective weight was given to the abundant ASVs, and the hill number decreased. The differences in microbiome compositions between sites (beta diversity) were estimated using both the pairwise Jaccard distance and Bray-Curtis dissimilarity index using the vegan package (Oksanen, 2020). Beta diversity was analyzed with non-metric multidimensional scaling (NMDS) and principal coordinates analysis (PCoA) using the phyloseq package (McMurdie and Holmes, 2013), and hierarchical clustering was performed with Ward’s method using the dendextend package (Galili, 2015). Pairwise permutational ANOVA (PERMANOVA) using the pairwiseAdonis package (Martinez Arbizu, 2017) was performed to test the differences in microbiome compositions between sites, with the Benjamini-Hochberg false discovery rate (FDR) correction based on 999 permutations, and the homogeneity of intra-group beta-dispersion was assessed using the phyloseq package (McMurdie and Holmes, 2013). The contribution of each ASV to the microbiome composition variation was assessed by similarity percentage (SIMPER) with the Bray-Curtis dissimilarity using the vegan package (Oksanen, 2020). Statistical significance was checked using the Kruskal-Wallis test by ranks. Only ASVs contributing more than 0.01% were retained.

To examine the effect of environmental and host factors on skin microbiome composition (beta diversity), two statistical models were used. In the first linear model, the relationship between geographical distance between sampling sites and microbiome similarity was tested. The pairwise Jaccard distance or Bray-Curtis dissimilarity was used as the response variable, and the geographical distance between sites was used as the explanatory variable. Pairwise distances between sites separated by the sea were excluded in the model, i.e., only pairwise distances between sites in the same region (New Territories, Hong Kong Island, or Lantau Island; Fig. 1b) were included. In the second linear mixed model, principal coordinate axis 1 (PCo1) was extracted from the Bray-Curtis dissimilarity or Jaccard distance matrix and used as the response variable. Elevation, BCI, and sex were included as fixed effects, and the sampling site and sampling month were included as random effects.

## Results

### Total, putative anti-*Bd*, and core skin microbiome of *P. hongkongensis*

Using high-throughput sequencing of the V4 region of the 16S rRNA gene, we identified 9,433 amplicon sequence variants (ASVs; Supplementary Table S4), belonging to 2,605 bacterial taxa classified to the lowest possible taxonomic level. In total, 42 phyla, 128 classes, and 274 orders of bacteria were identified. Among the identified ASVs, 314 were found in the putative anti-*Bd* bacteria database (Woodhams et al., 2015), assigned to 25 genera (Supplementary Table S5), accounting for 9.7% of the total reads. On average, each *P. hongkongensis* individual harbored 21 putative anti-*Bd* ASVs. All individuals harbored at least three putative anti-*Bd* ASVs. The most prevalent putative anti-*Bd* ASV was present in 86.6% of all individuals, and 114 putative anti-*Bd* ASVs were uniquely found in single individuals. Two ASVs from *Acinetobacter*, three from *Flavobacterium* and three from *Novosphingobium* had the highest relative read abundance (RRA) and occurrence. Rarefaction indicated sufficient sequencing depth for ASV identification (Supplementary Fig. S1).

The most abundant phyla were Proteobacteria (59.2%), Bacteroidetes (21.2%), and Verrucomicrobia (12%), together making up more than 92% of the total reads, while no other phyla had an RRA >1% (Fig. 2). The most abundant classes in Proteobacteria were Betaproteobacteria (44.8%), Gammaproteobacteria (6.5%), Alphaproteobacteria (6.4%), Cytophagia (1.2%), and Deltaproteobacteria (1%), while the most abundant among Bacteroidetes were Sphingobacteriia (5.9%) and Flavobacteria (3.5%), and most abundant in Verrucomicrobia were Opitutae (6.2 %) and Verrucomicrobiae (5.7%). Comamonadaceae, a family in Betaproteobacteria, was the most abundant taxon, represented by 33.3% of the total reads, and present in every individual skin microbiome (Fig. 2). Five out of 16 ASVs, each accounting for >1% reads, were assigned to Comamonadaceae (Supplementary Table S6). There were 14 taxa with an RRA >1%, among which three were putatively anti-*Bd* (*Acinetobacter*, *Flavobacterium*, and *Novosphingobium*; Supplementary Table S5). The core microbiome comprised 13 bacterial taxa (Supplementary Table S7) represented by 7.4% of the total reads, two of which were putatively anti-*Bd* (*Flavobacterium* and *Novosphingobium*; Supplementary Table S5). All taxa that accounted for >1% of the total reads or were within the core microbiome belonged to one of the three most abundant phyla. In addition, ASVs from potentially TTX-producing *Aeromonas*, *Pseudomonas*, *Shewanella*, and *Sphingopyxis* were found in our dataset, but only ASV21 (*Aeromonas* sp.) was present in a majority of newt individuals (60%) and water samples (75%), with its RRA highly variable across samples (mean=0.4%, S.D.=1.68%, range=0-13.9%; Supplementary Tables S8, S9).

**Figure 2.**
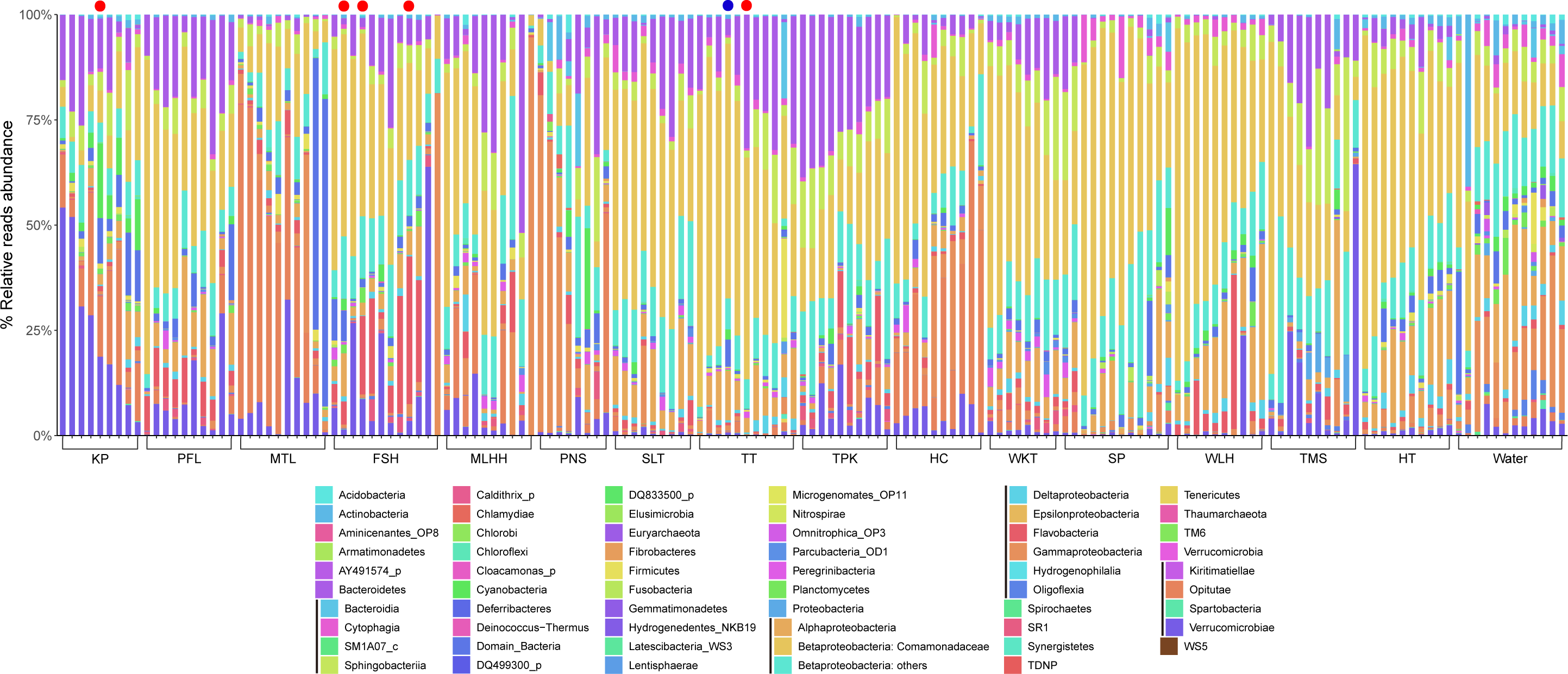
Skin microbiome compositions of *P. hongkongensis* by sampling sites. Relative read abundances of bacterial taxa of each skin swab sample are shown. All taxa are shown at the phylum level except that the three most abundant phyla, Proteobacteria, Bacteroidetes, and Verrucomicrobia, are reported at the class level. Color-coded taxon in columns from left to right are presented in a top-down manner in each bar. The most abundant taxon, the family Comamonadaceae (phylum: Proteobacteria; class: Betaproteobacteria), is specifically indicated. *Bsal* and *Bd* positive samples are marked by a red dot and a blue dot, respectively, above the corresponding bar. Refer to Fig. 1 for the abbreviations of sampling sites.

### Microbiome diversity within and between sites

Microbiome composition and diversity differed among sites and between skin and water samples (Figs. 2-3; Supplementary Fig. S2; Supplementary Tables S10, S11, S12). Individual microbiomes from Hok Tau (HT), Kowloon Peak (KP), and Wu Kau Tang (WKT) had the highest ASV richness (at *q* = 0), while those from Tai Mo Shan (TMS) had the lowest (Figs. 1b, 3a; Supplementary Tables S13, S14). HT and KP also showed the highest gamma diversity in microbial communities (Fig. 3b). As *q* increased, both the alpha diversity (within each sample) and gamma diversity (within each site) of the newt skin microbiomes from all sites decreased rapidly, reflecting that the relative abundances of bacterial taxa in the skin microbiomes were highly uneven.

**Figure 3.**
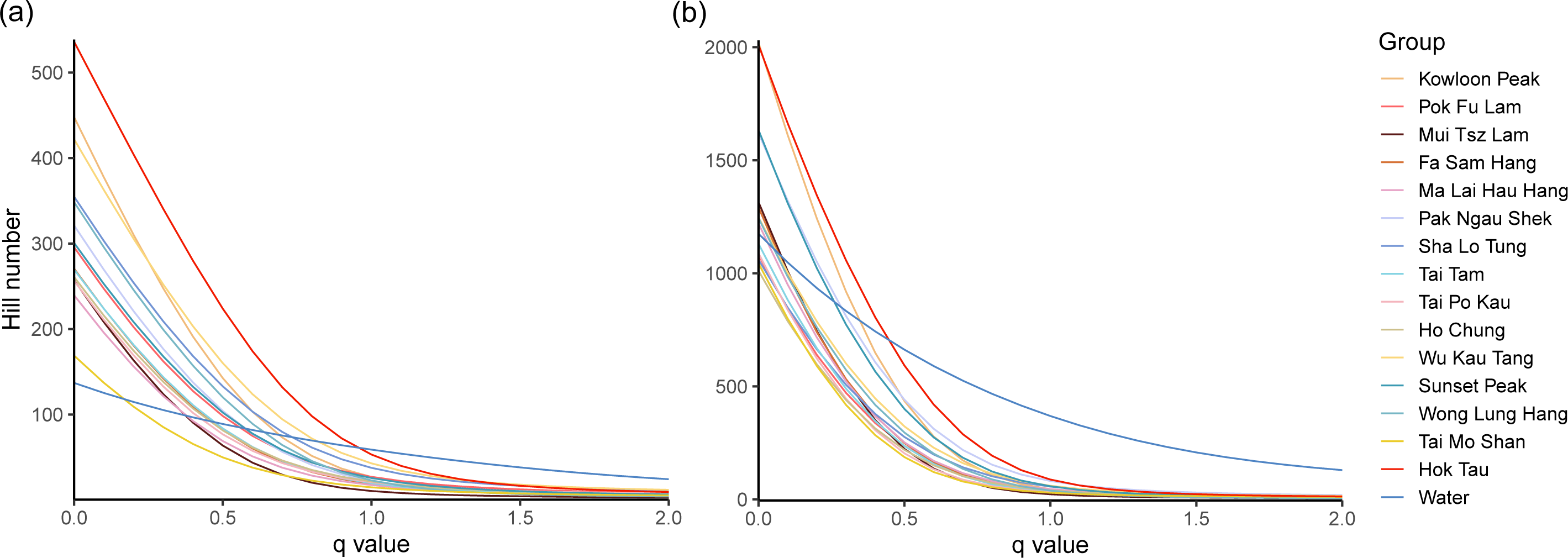
Alpha and gamma diversity of *P. hongkongensis* skin microbiomes from 15 sampling sites. (a) Alpha diversity - the mean effective number of ASVs in an individual sample for each site and (b) Gamma diversity - the total effective number of ASVs found in each site, when *q* ranges from 0 to 2 are shown.

Microbiome composition analyses showed that the skin microbiomes demonstrated site differences and were differentiated from water microbiomes (hierarchical clustering, Fig. 4; non-metric multidimensional scaling [NMDS] and principal coordinates analysis [PCoA], Supplementary Fig. S3). Skin microbiomes were well differentiated by site in the hierarchical clustering analysis based on both the Bray-Curtis dissimilarity (Fig. 4; Supplementary Table S15) and Jaccard distance (Supplementary Fig. S4; Supplementary Table S16), indicating site-specific bacterial compositions. The Bray-Curtis dissimilarity analyses revealed that skin microbiomes from Sunset Peak (SP) and Wong Lung Hang (WLH) were highly similar to each other (Figs. 1b, 4). TMS had a relatively distinct skin microbiome composition from most sites according to the results from NMDS (Supplementary Fig. S3a) and PCoA (Supplementary Fig. S3c; Supplementary Table S17). Jaccard distance analyses presented similar patterns to the Bray-Curtis dissimilarity results in NMDS (Supplementary Fig. S3b) and PCoA (Supplementary Fig. S3d; Supplementary Table S18). Bacterial communities in water samples were clustered as a distinct group in Bray-Curtis dissimilarity analyses, except the one from SP which grouped with SP skin microbiome samples (Fig. 4; Supplementary Figs. S3, S4).

**Figure 4.**
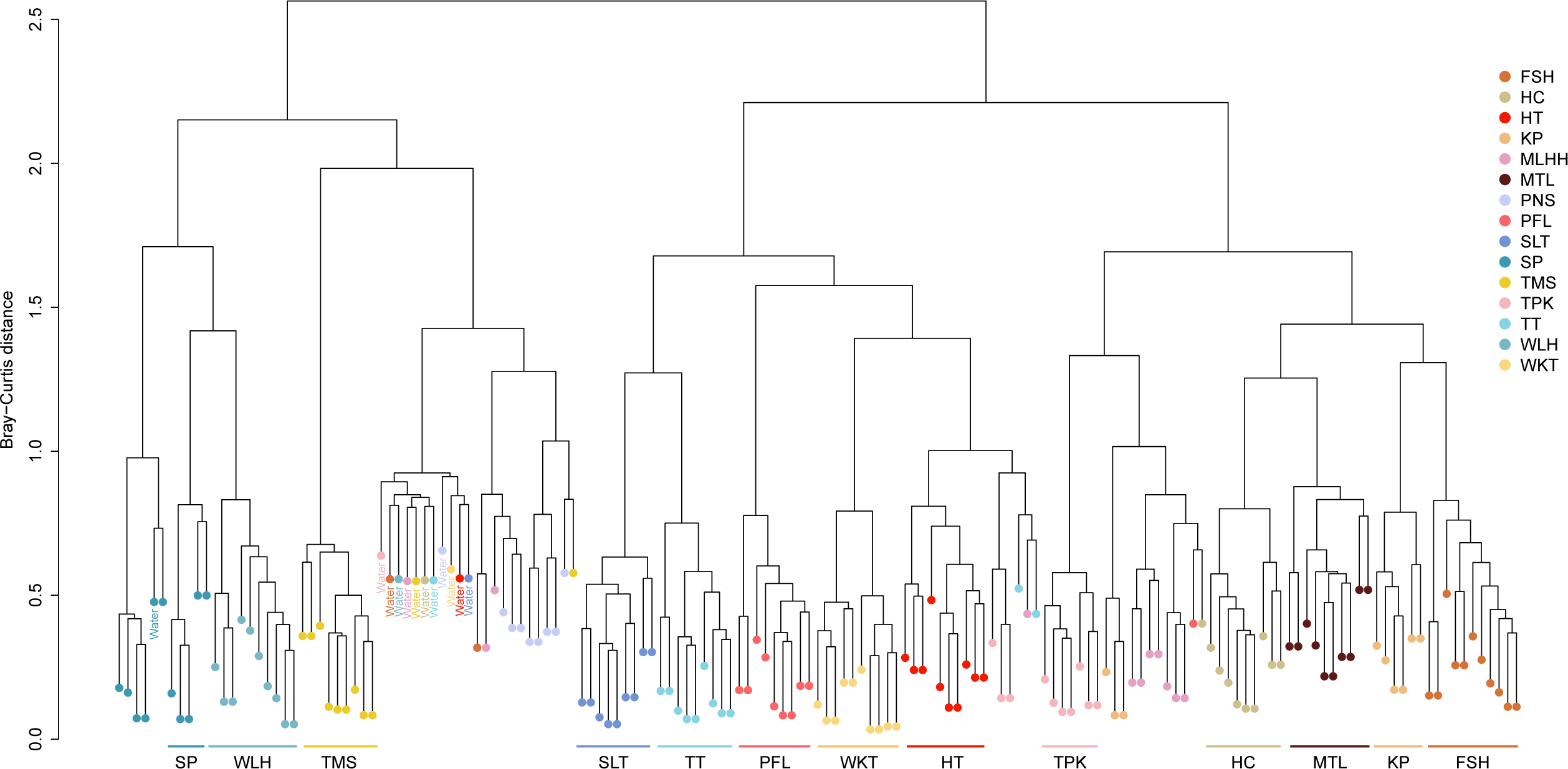
Hierarchical clustering of *P. hongkongensis* skin microbiomes by sampling sites based on the Bray-Curtis dissimilarity. Clusters with at least five samples from the same site are highlighted by the site name. Refer to Fig. 1 for the abbreviations of sampling sites.

The skin microbiome composition was significantly different between all pairwise site comparisons (adjusted *p* ≤ 0.002; Supplementary Tables S19, S20). The mean number of ASVs contributing to the top 10% dissimilarity between any two sites was 30.1 (S.D. = 8.7). Within the top 10% dissimilarity, seven ASVs were present in more than 50% of the 105 pairwise comparisons (Supplementary Table S21), while 90 ASVs contributed to the dissimilarity of just one or two site pairs. Site-specific patterns were observed in the composition of newt skin microbiomes (Figs. 2, 4; Supplementary Fig. S4). While the bacterial communities from most sites were dominated by Betaproteobacteria, those from Mui Tsz Lam (MTL) had a large proportion of Opitutae (39.3% RRA; Fig. 2). Verrucomicrobiae was the most abundant class among the samples from KP (25.8% RRA; Fig. 2). In half of the samples from Pak Ngau Shek (PNS), Gammaproteobacteria were present in the highest abundance (58% RRA; Fig. 2). Skin microbiomes from SP and WLH, the two sites on Lantau Island, as well as TMS, the highest peak in Hong Kong (Fig. 1), were each represented by relatively well-separated groups in NMDS (Supplementary Fig. S3a, b). Compared to other sites, bacterial communities from MTL, SP, and WLH had a particularly low abundance of Bacteroidetes (Fig. 2). Microbiomes in water samples appeared more evenly distributed across taxa than skin microbiomes (Fig. 3) and were more similar to each other than to skin microbiomes (Fig. 4; Supplementary Figs. S3, S4).

### Effects of environmental and host factors on skin microbiome

The pairwise distance between microbiomes was positively correlated to the geographical distance between sites in terms of Bray-Curtis dissimilarity (*p* < 0.001, *R*^2^ = 0.21; Fig. 5a) and Jaccard distance (*p* < 0.0001, *R*^2^ = 0.23; Fig. 5b). The linear mixed model (Table 1) suggested that the skin microbiome composition changed with site elevation (*p* = 0.02; Fig. 5c) and was significantly different between host sexes (*p* < 0.01; Fig. 5d).

**Figure 5.**
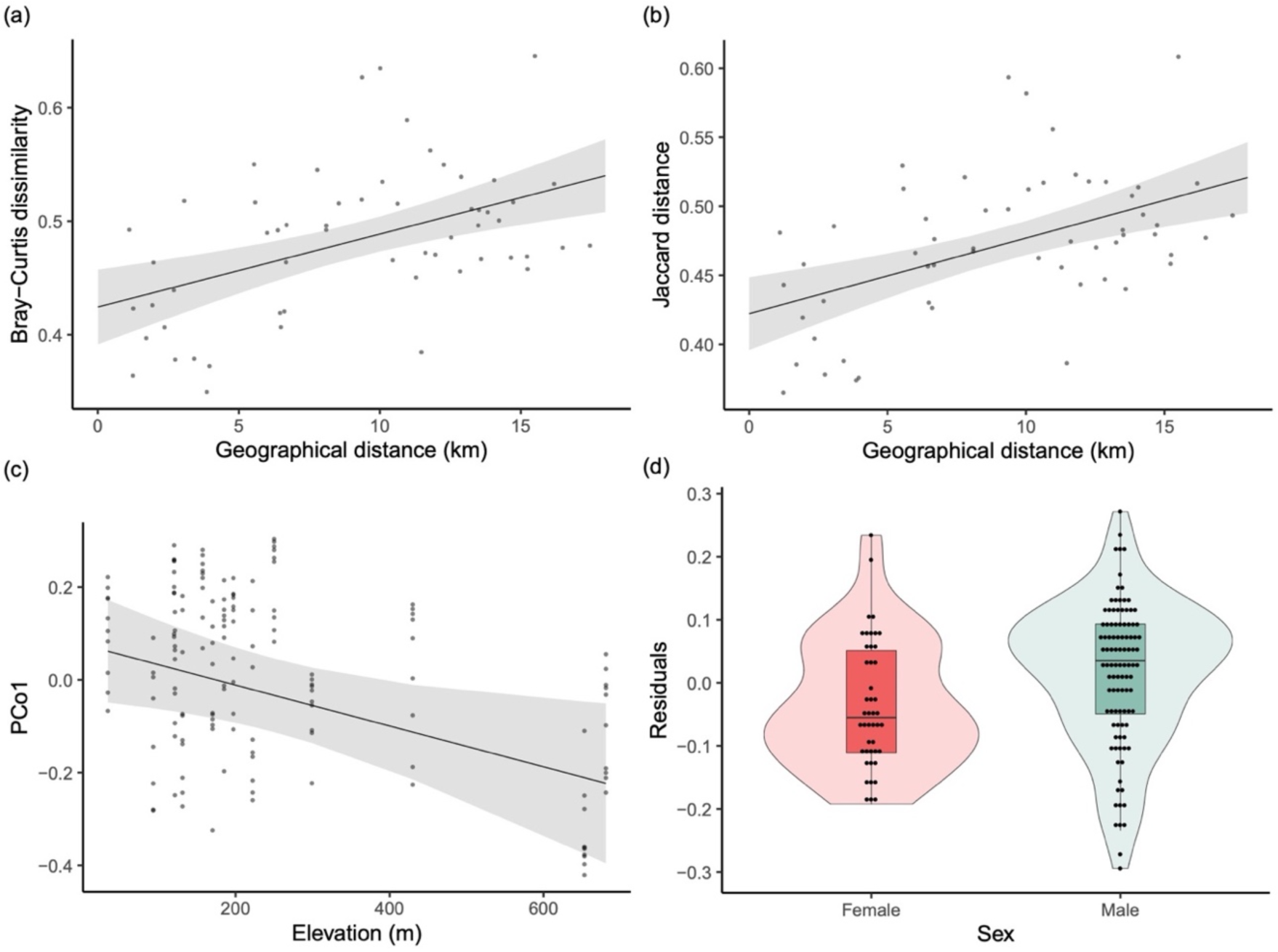
Effects of environmental and host factors on *P. hongkongensis* skin microbiome. The (a) Bray-Curtis dissimilarity (*p*<0.001, *R*^2^=0.21) and (b) Jaccard distance (*p*<0.0001, *R*^2^=0.23) between the microbiomes from different sites increased with the geographical distance between sites. (c) Microbiome composition (represented by PCo1 based on Bray-Curtis dissimilarity) varied with site elevation (*p*=0.02). (d) The difference in microbiome composition between sexes (*p*<0.01), shown as the differential residual distributions in the fitted model where sex was not included.

**Table 1.**
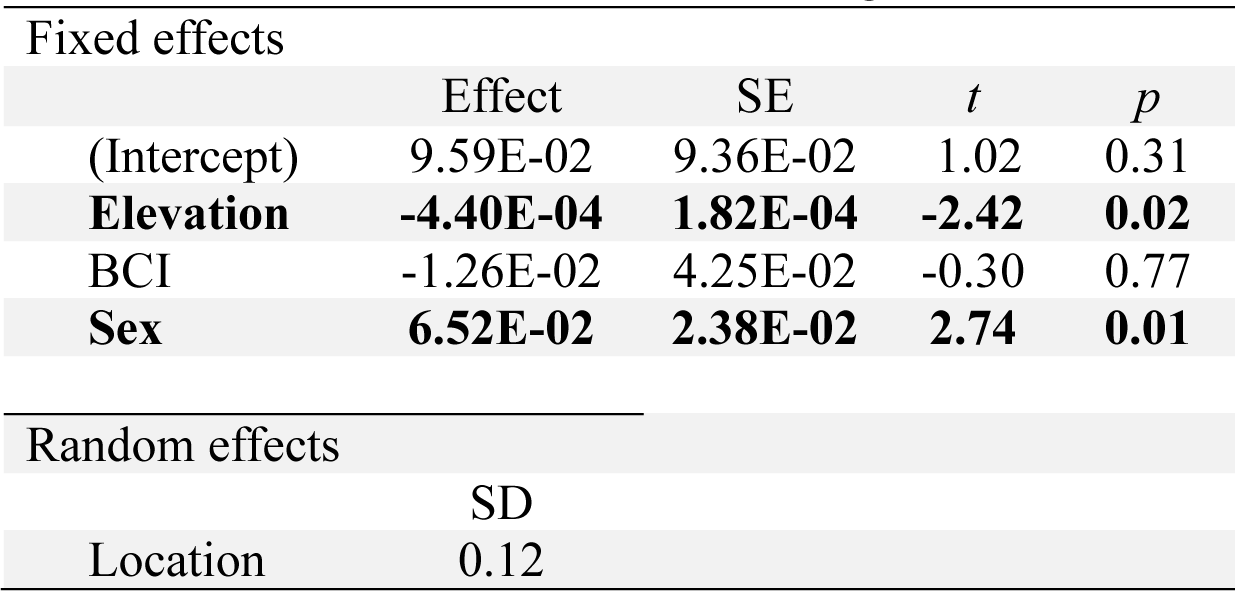
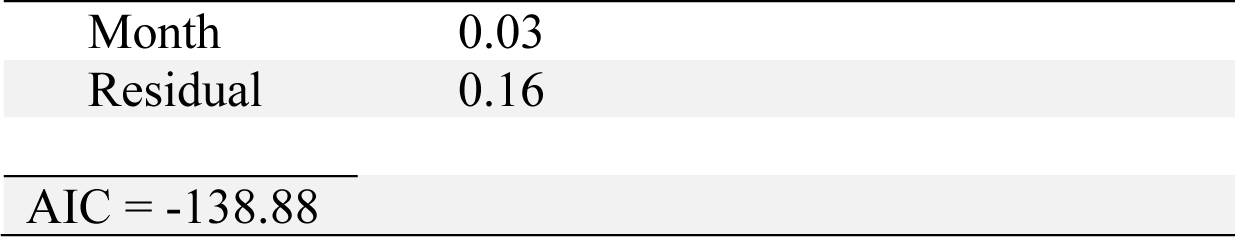
Linear mixed model assessing the effects of environmental and host factors on the skin microbiome of *P. hongkongensis* (n=147). Site elevation, body condition index (BCI), and sex were included as fixed effects. Sampling location and month were included as random effects. Significant effects are bolded.

| Fixed effects |  |  |  |  |
| --- | --- | --- | --- | --- |
|  | Effect | SE | *t* | *p* |
| (Intercept) | 9.59E-02 | 9.36E-02 | 1.02 | 0.31 |
| **Elevation** | **-4.40E-04** | **1.82E-04** | **-2.42** | **0.02** |
| BCI | -1.26E-02 | 4.25E-02 | -0.30 | 0.77 |
| **Sex** | **6.52E-02** | **2.38E-02** | **2.74** | **0.01** |
| Random effects |  |  |  |  |
|  | SD |  |  |  |
| Location | 0.12 |  |  |  |

|  |  |
| --- | --- |
| Month | 0.03 |
| Residual | 0.16 |
| AIC = -138.88 |  |

## Discussion

As the pandemic caused by chytrid fungi continues to threaten global amphibian diversity, Asia is gaining more attention from the scientific community after being inferred as the origin of the pathogens (Fisher and Garner, 2020, O’hanlon et al., 2018). This study is one of the few to investigate the skin microbiome in wild amphibian populations in Asia (Bletz et al., 2017b, Sabino-Pinto et al., 2016, Mutnale et al., 2021, Bataille et al., 2018, Yang et al., 2020). We found that the diverse bacterial communities on the skin of *P. hongkongensis* were dominated by Proteobacteria, Bacteroidetes, and Verrucomicrobia and contained a great diversity of putative anti-*Bd* bacteria. The compositions of the skin microbiomes were significantly different across sampling sites, and most of them were highly differentiated from the stream water microbiomes present in the habitats of *P. hongkongensis*. The dissimilarity between skin microbial communities was found to be correlated with the geographical distance between sites. Moreover, among other environmental and host factors tested, elevation and host sex were also identified to be significant factors in determining the skin microbiome structure of the newts.

In congruence with many existing skin microbiome studies in amphibians, Proteobacteria was the most abundant bacterial phylum on *P. hongkongensis*, followed by Bacteroidetes. Comamonadaceae was also commonly identified as an abundant family (Belden et al., 2015, McKenzie et al., 2012, Bletz et al., 2017a, Jiménez et al., 2019, Prado-Irwin et al., 2017, Walker et al., 2020, Walke et al., 2021, Bletz et al., 2018). While Verrucomicrobia was occasionally found among the dominant phyla, its relative abundance seldom exceeded 3% (Belden et al., 2015, Walke et al., 2021, Bletz et al., 2018, Bletz et al., 2017b), with exceptions in the Japanese giant salamander (*Andrias japonicus*, 9%) and the larvae of wild Japanese fire belly newts (*Cynops pyrrhogaster*, 20%) inhabiting Japan (Sabino-Pinto et al., 2016). Yet a few phyla that are frequently found on most amphibian skin with high abundance, such as Actinobacteria, Firmicutes, Acidobacteria, and Cyanobacteria (Belden et al., 2015, McKenzie et al., 2012, Bletz et al., 2017a, Jiménez et al., 2019, Prado-Irwin et al., 2017, Walker et al., 2020, Walke et al., 2021, Bletz et al., 2018), were scarce (RRA< 1%) on *P. hongkongensis*. In a study on the skin microbial community structures of 89 frog species in Madagascar, Actinobacteria and Acidobacteria were shown to occur in significantly higher relative abundance on terrestrial species than on aquatic or semiaquatic species, which might explain their low abundance on *P. hongkongensis* which is mostly aquatic during the sampling period (Bletz et al., 2017a).

Although there were large intraspecific variations in the skin microbiomes across sites, a group of core microbiome members were shared by most *P. hongkongensis* individuals, even though their relative abundances were generally low. The maintenance of a core microbiome, which displays a relatively stable composition under different environmental conditions experienced by the hosts, was observed in many amphibian species, suggesting its functional importance. For example, the structures of the core microbiome on coqui frogs (*Eleutherodactylus coqui*) did not vary along the gradient of elevation or forest integrity (Hughey et al., 2017). The core microbiome of *P. hongkongensis* comprised of bacterial species that were likely well adapted to reside on the host skin and performing vital functions for the host, resulting in a type of symbiosis (Prado-Irwin et al., 2017, Muletz Wolz et al., 2018). However, since we only sampled during the breeding season of the newts, the abundance of these core bacterial species may fluctuate temporally as the newts move from an aquatic to a terrestrial environment after the breeding season, causing changes to a number of relevant host traits, e.g., the levels of skin moisture or reproductive hormones (Bletz et al., 2017b, Lau et al., 2017, Lau, 2017). Seasonal changes in the abundance of core microbes, e.g., Chitinophagaceae, a core family also detected on *P. hongkongensis*, has been reported in the North American wood frog (*Rana sylvatica*) (Douglas et al., 2021). Noteworthily, *P. hongkongensis* produce TTX (Yotsu et al., 1990), which was proposed to act as a defense against predators, but the synthesis mechanism of this deadly neurotoxin in *P. hongkongensis* is unknown. Further investigation of the TTX-producing ability of the *Aeromonas* sp. found on many newt individuals can inform us if *P. hongkongensis* has developed a symbiotic relationship with certain toxin-producing bacteria to increase its fitness, like in another poisonous newt (Vaelli et al., 2020).

Furthermore, core microbiome might be functionally important in boosting host resistance against *Bd* infection. For example, based on bioinformatic and culture tests, it was predicted that *P. cinereus* has maintained a core microbiome in which bacterial members similarly provided anti-*Bd* functions (Loudon et al., 2014). Multiple species of *Novosphingobium* and *Flavobacterium* detected as the core microbes on *P. hongkongensis* have already been shown to exhibit anti-*Bd* property (Harris et al., 2006), with the latter genus being widely distributed among amphibian species, such as *R. cascadae* (Roth et al., 2013), the green-eyed frog (*Lithobates vibicarius*) (Jiménez et al., 2019), and the four-toed salamander (*Hemidactylium scutatum*) (Lauer et al., 2008). Comamonadaceae, *Methylotenera*, and *Undibacterium* were also identified as the core taxa on the wild Oriental fire-bellied toad (*Bombina orientalis*) in Japan (Sabino-Pinto et al., 2016), and the abundance of *Methylotenera* and Rhizobiales on wild *B. orientalis* was found to vary with individual *Bd* infection status in South Korea (Bataille et al., 2016, Bataille et al., 2018). In general, although the functions of core microbiomes on amphibian skin are not well understood, the overlapping skin microbiome composition between *P. hongkongensis* and other Asian amphibians studied hints at a potential protective role of their core microbes against chytrid fungal pathogens.

It may be reasonable to assume the putative anti-*Bd* bacterial taxa identified here using the anti-*Bd* isolates database could potentially possess anti-*Bsal* property as well, since a lot of these bacteria can produce broad antifungal compounds (Woodhams et al., 2015, Muletz Wolz et al., 2018). Putative antifungal bacteria were detected on *P. hongkongensis* in great diversity, high prevalence, but low abundance. Eight ASVs from *Acinetobacter*, *Flavobacterium* and *Novosphingobium* were particularly abundant and prevalent, suggesting their putative symbiotic relationship with *P. hongkongensis* (Muletz-Wolz et al., 2017). Most putative anti-*Bd* microbiomes had restricted distributions in newt individuals and dissimilar taxon identities, which might have resulted from the colonization by putative anti-*Bd* bacteria acquired from distinct pools of microbes present in different locations (Muletz-Wolz et al., 2017). The estimated relative abundances of the putative anti-*Bd* bacteria in this study were generally much lower than those found in some other amphibian species, such as three *Plethodon* species in North America (Muletz Wolz et al., 2018) and six frog species in India (Mutnale et al., 2021). This might reflect the functional importance of rare taxa in a microbiome. Another explanation could be the failure to capture significant antifungal members present in the microbiomes of *P. hongkongensis* via identification using the anti-*Bd* bacteria database. Since the database has been compiled by testing bacterial isolates from certain amphibian species with culture methods, rare or unculturable bacteria and host-specific microbial species on untested amphibians, especially those in Asia, are therefore unlikely to be included. Nonetheless, the fact that putative anti-*Bd* bacteria were detected in all 100% newts sampled indicated the potential resistance or even herd immunity against chytridiomycosis in *P. hongkongensis*, possibly a result of historical interactions with *Bd* and *Bsal* strains endemic to Asia, where *Bd* has a high degree of diversity and has not been found to cause morbidity or mortality in local wild amphibian populations (Bataille et al., 2013).

It has been suggested that an amphibian population can achieve herd immunity to *Bd* when around 80% of the individuals are protected by anti-*Bd* microbes, enabling the population to coexist with *Bd* (Woodhams et al., 2007, Muletz-Wolz et al., 2017). In the case of *P. hongkongensis* population in Wutongshan, all *Bsal*-positive individuals showed no symptoms despite the high prevalence of the pathogen (Yuan et al., 2018). For the *P. hongkongensis* population in Hong Kong, the prevalence of *Bd* and *Bsal* was very low (Chen et al., 2023) with positive individuals being asymptomatic, supporting the idea of potential herd immunity achieved through antifungal skin microbes. In this study, no noticeable difference was observed in the skin microbiomes between *Bsal*-positive and *Bsal*-negative individuals. Noteworthily, *Janthinobacterium* spp. (Class Betaproteobacteria, Phylum Proteobacteria) were detected on *P. hongkongensis*. The anti-*Bd* property of *Janthinobacterium lividum* was well verified (Brucker et al., 2008). This bacterium has been shown to produce indole-3-carboxaldehyde and violacein that could significantly inhibit *Bd* growth at low concentrations (Brucker et al., 2008), and its addition to amphibian skin could reduce both the morbidity and mortality caused by *Bd* infection (Becker et al., 2009, Harris et al., 2009a). In addition, there were also numerous ASVs found on *P. hongkongensis* that belonged to a few common anti-*Bd* genera such as *Pedobacter* and *Pseudomonas*.

One of our key findings was that the skin microbiome structure of newts was largely differentiated by site, and was dependent on environmental or host genetic factors. The population genetic structure of *P. hongkongensis* has been previously shown to accord with the landscape topography, which was poorly explained by isolation-by-distance (Lau, 2017). Based on the population structure, there were five genetic clusters of *P. hongkonggensis*, one on Hong Kong Island [Pok Fu Lam (PFL), Tai Tam (TT)], one on Lantau Island (SP), and three in the New Territories [North: WKT; East: HC, MTL, KP; and Central: Tai Po Kau (TPK)], while an admixture of the New Territories North and East clusters was observed at PNS (Lau, 2017). Consistent with the host population structure, the skin microbiomes from the two sites (WLH and SP) on Lantau Island were similar to each other and largely different from other sites (Fig. 4; Supplementary Fig. S4). The geographical isolation of the Lantau Island populations from the mainland not only limited the gene flow between newts, but probably also the exchange of individual microbiomes. However, this pattern was not observed between the two Hong Kong Island newt populations (PFL and TT), where their skin microbiomes were more similar to those of the New Territories populations than to each other, implying other factors might also influence microbiome structure besides geographical isolation alone. Microbiome differences in the New Territories can be explained in part by geographical and host genetic distances, e.g., individuals from the New Territories East sites (i.e., HC, MTL, KP, and FSH) shared mostly similar microbiomes, but they could not account for all differences. Hence the microbiomes from some sites were not grouped according to host population structure or geographical proximity. Overall, geographical distance appeared to play a larger role since within-site microbiomes strongly clustered together but not intermixed with others from the same genetic population of newts from a different site (Fig. 4; Supplementary Fig. S4).

How amphibians acquire their skin microbes in each new generation and how their microbiomes are shaped and maintained remain important questions in the field.

Horizontal transmission happens when microbes are selected from the habitat to form a microbiome (Bright and Bulgheresi, 2010). Members of the *P. hongkongensis* skin microbiome might come from the living environment, such as water, soil, and leaf litter (Muletz Wolz et al., 2018). The clear difference in the microbiomes between skin and stream water implies that even though the skin of *P. hongkongensis* might initially contact all bacterial species present in the surrounding, only a subset of these species had managed to colonize and proliferate on the skin. Since the environmental conditions, e.g., water quality (Krynak et al., 2016, Kueneman et al., 2014), soil, and pH (Muletz Wolz et al., 2018, Fierer and Jackson, 2006), of various sites could be different, the composition of bacteria available from the environment for horizontal transmission could certainly vary in response to the site-specific conditions. Moreover, the environmental characteristics at each site could affect host traits, so the site-wise differences in microbiomes could also reflect the influence of site-specific host traits.

For instance, the production of antimicrobial peptides by the skin, a key component in amphibian immune defense, could be regulated by water surface area and conductivity, thus affecting the skin microbiome composition (Krynak et al., 2016).

In addition to geographical distance, elevation also had an effect on the skin microbial communities of newts, similar to previous observations that skin microbiome structures varied along elevational gradients from 0 to 875 m and 700 to 1,000 m above sea level (Muletz Wolz et al., 2018, Hughey et al., 2017). The influence of elevation is complicated and many environmental factors co-vary along the gradient, which might in turn affect the host traits as well as the growth and interactions of different bacteria on the host (Muletz Wolz et al., 2018). Furthermore, *Bd* prevalence was found to positively correlate with elevation, which might have led to a stronger selection for anti-*Bd* bacteria on surviving hosts at higher elevations if there have been recurring infections (Bresciano et al., 2015).

We also found that the variations in the microbiome structures of *P. hongkongensis* were associated with host sex. Although it has been shown in Blanchard’s cricket frog (*Acris blanchardi*) that sex interacted with latitude and water surface area in affecting the skin microbiome (Krynak et al., 2016), it has no effect on *E. eschscholtzii* (Prado-Irwin et al., 2017) or *R. sylvatica* (Douglas et al., 2021). In *P. hongkongensis*, females exhibited cannibalism on eggs and larvae, which caused a difference in the diets between sexes (Fu et al., 2013). As diet can influence the skin microbiome (Antwis et al., 2014), the effect of sex might, in fact, be an indirect indicator of the effect of diet. Alternatively, hormonal differences between the sexes might explain their microbiome divergence. A relationship between sexual differences in host-associated microbiomes and sex hormone levels has been documented in birds and primates (Escallon Herkrath, 2015, Mallott et al., 2020), but whether such a relationship exists in amphibian skin microbiomes has not been investigated.

The skin microbiome of *P. hongkongensis* was diverse, with a group of core skin microbes potentially coevolving with, and providing essential functions for, the hosts. The bacterial community structure on *P. hongkongensis* was highly site-specific, which was likely attributed to geographical isolation and environmental characteristics instead of host population genetics, as demonstrated by the effect of elevation. Host sex also influenced the newt skin microbiome, possibly because of the differences in diet or hormonal levels between the two sexes. The high prevalence and low abundance of putative antifungal bacteria found on the skin, together with the low prevalence and dispersed distribution of chytrid infection in the newt population, suggest that *P. hongkongensis* has had a history of exposure to and coexistence with the fungal pathogens in the region. Using *P. hongkongensis* as a reference species, interrogation of the compositions and patterns of the skin microbiome structures of other amphibians in East Asia, where chytrids are proposed to be native to, will enhance our understanding on host-pathogen-microbiome coevolution. Such knowledge can be applied to the conservation of *P. hongkongensis* as well as amphibian populations naïve to the pathogenic chytrids. Given that the extant *P. hongkongensis* skin microbiome has likely been shaped by the coevolution with chytrids, it is a promising source for identifying and characterizing antifungal bacteria not yet known to us, and thereby developing novel probiotic treatments against chytridiomycosis. Further investigations into the factors governing the differences in the skin microbiomes between sites and sexes could improve our understanding and our ability to develop effective probiotic treatments.

## Data accessibility

The datasets generated for this study will be deposited to NCBI. (Accession numbers will be provided after the manuscript is accepted.)

## Author contributions

B.W.: Investigation, Formal analysis, Visualization, Writing - Original Draft; G.C.: Investigation, Formal analysis, Writing - Review & Editing; E.S.K.P.: Investigation, Methodology, Writing - Original Draft, Writing - Review & Editing; H.S.F.: Investigation; A.L.: Investigation; S.Y.W.S.: Conceptualization, Investigation, Writing - Review & Editing, Resources, Supervision, Funding acquisition, Project administration.

## Acknowledgments

We would like to thank Yik-Hei Sung and Wing Ho Lee for their assistance during sample collection, Pei-Yu Huang for her advice on analysis, and Charis May Ngor Chan for her technical assistance. The computations were performed using the research computing facilities offered by the Information Technology Services at the University of Hong Kong. This work was supported by the Start-Up Fund granted to S.Y.W.S. by the University of Hong Kong. This work was approved by the Agriculture, Fisheries and Conservation Department [Permit no.: (29) in AF GR CON 09/51Pt.7] of the HKSAR Government.

---

**Figure S1.**
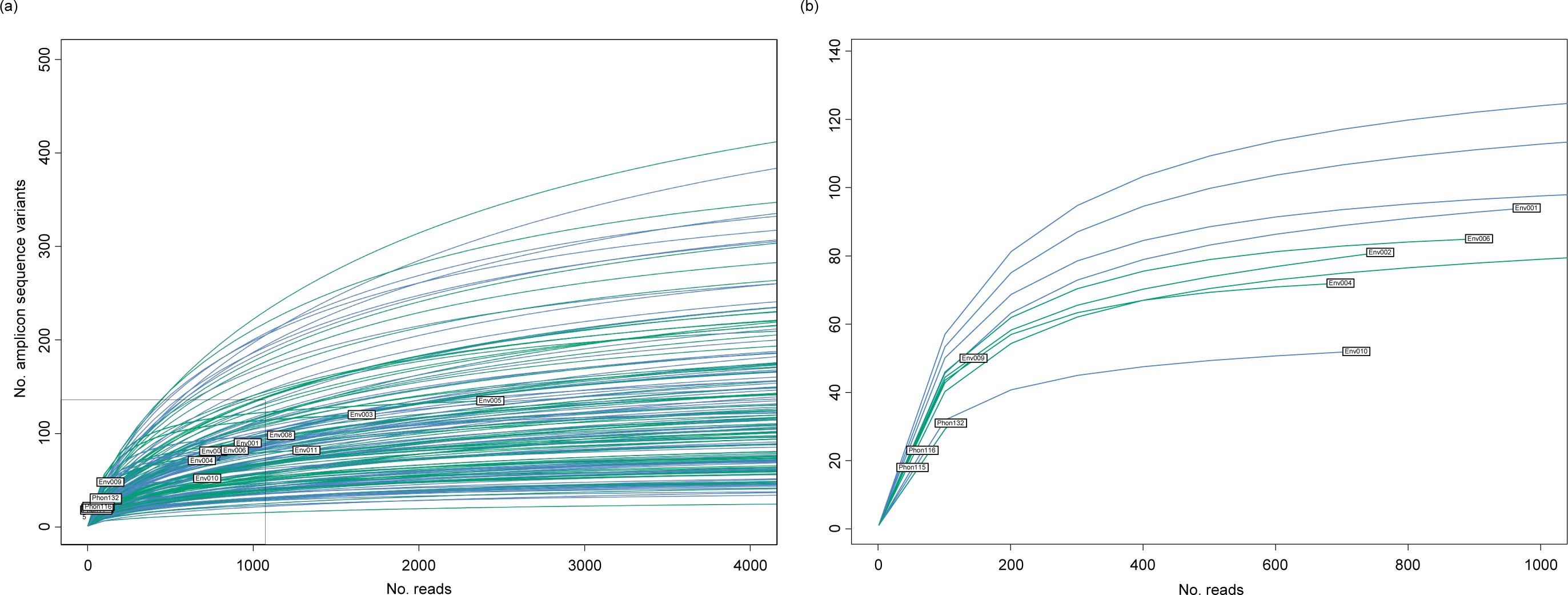
Rarefaction curves of all skin swab samples of *P. hongkongensis*. (a) Most samples have a plateauing rarefaction curve, which indicates sufficient sequencing depths. (b) Enlarged view of the indicated area in (a) showing samples with the lowest numbers of reads. Phon115, Phon116, and Phon132 did not have sufficient reads and were therefore excluded from subsequent analyses.

**Figure S2.**
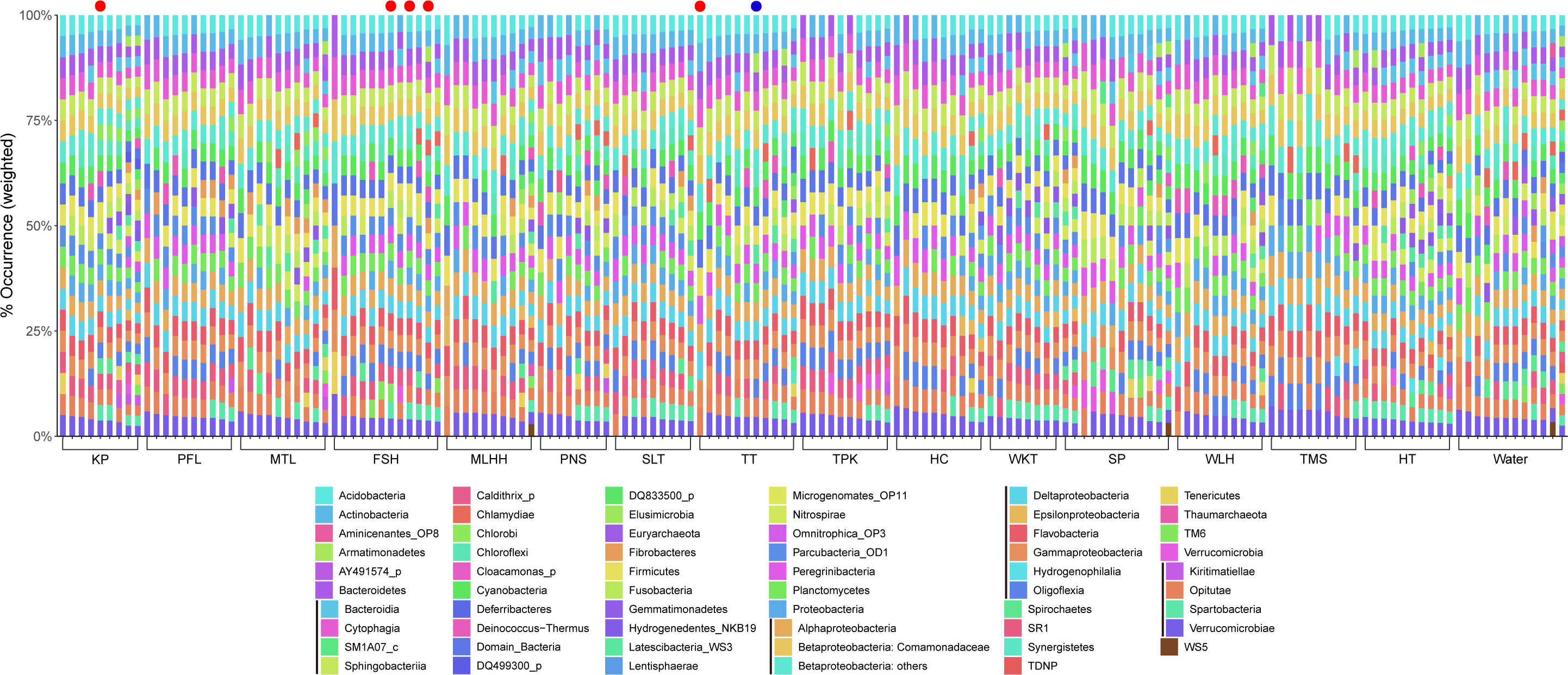
Skin microbiome composition of *P. hongkongensis* by sampling sites shown in weighted percentage of occurrence. All taxa are shown at the phylum level except that the three most abundant phyla, Proteobacteria, Bacteroidetes, and Verrucomicrobia are reported at the class level. Color-coded taxon in columns from left to right are presented in a top-down manner in each bar. The most abundant taxon, the family Comamonadaceae (phylum: Proteobacteria; class: Betaproteobacteria) is specifically indicated. *Bsal* and *Bd* positive samples are marked by a red dot and a blue dot above the corresponding bar, respectively. Refer to Fig. 1 for the abbreviations of sampling sites.

**Figure S3.**
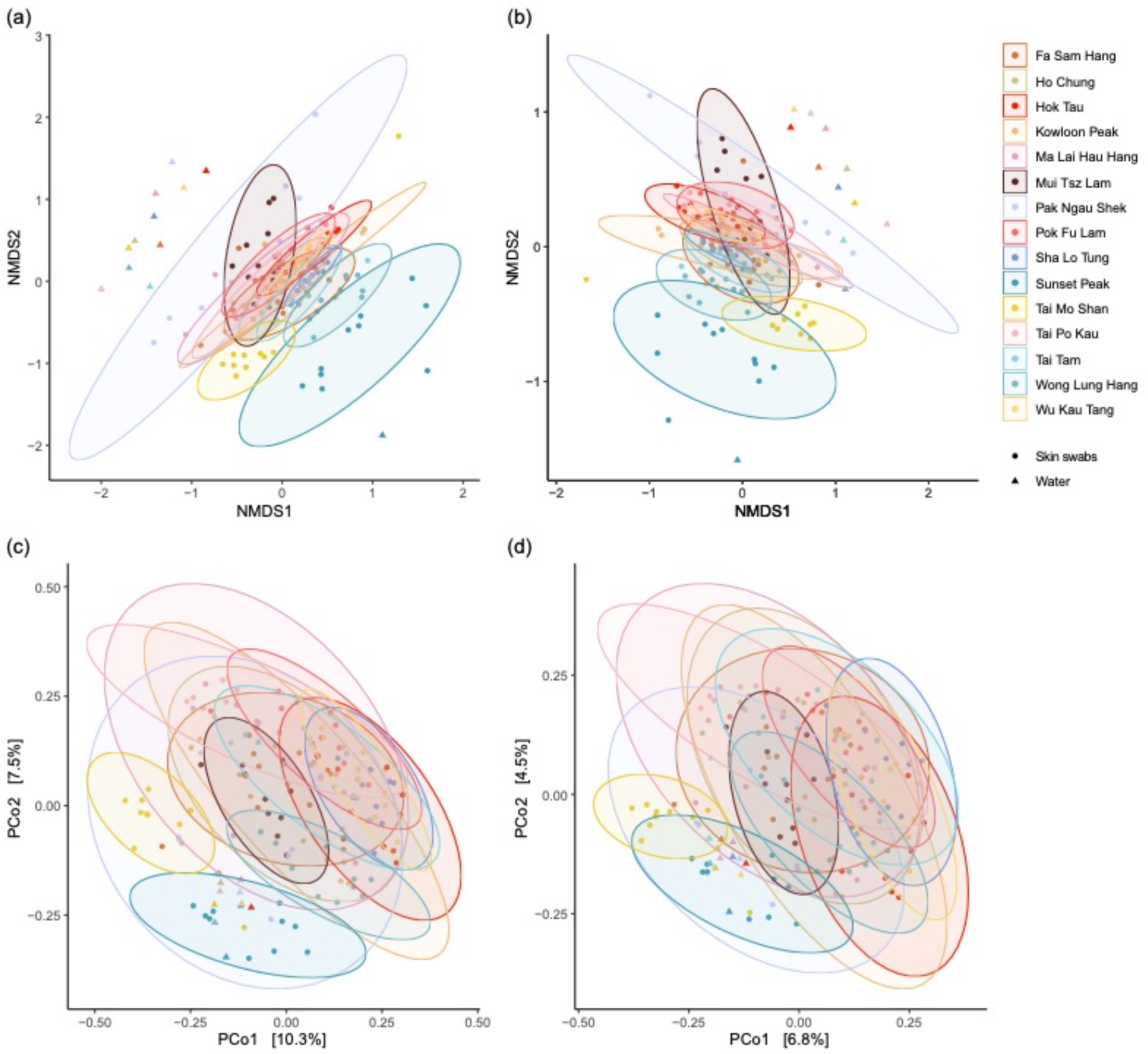
Non-metric multidimensional scaling (NMDS) and principal coordinates analysis (PCoA) of *P. hongkongensis* skin microbiomes from 15 sampling sites. NMDS plots based on the (a) Bray-Curtis dissimilarity and (b) Jaccard distance similarly demonstrate that *P. hongkongensis* skin microbiomes from Sunset Peak (SP), Wong Lung Hang (WLH), and Tai Mo Shan (TMS) formed groups relatively distant from those from other sites. Water microbiomes showed distinct composition compared to skin microbiomes. Pattens in the PCoA plots based on the (c) Bray-Curtis dissimilarity and (b) Jaccard distance appear highly similar. In accordance with NMDS, the skin microbiomes from SP and TMS are dissimilar to others. Water microbiomes form a cluster.

**Figure S4.**
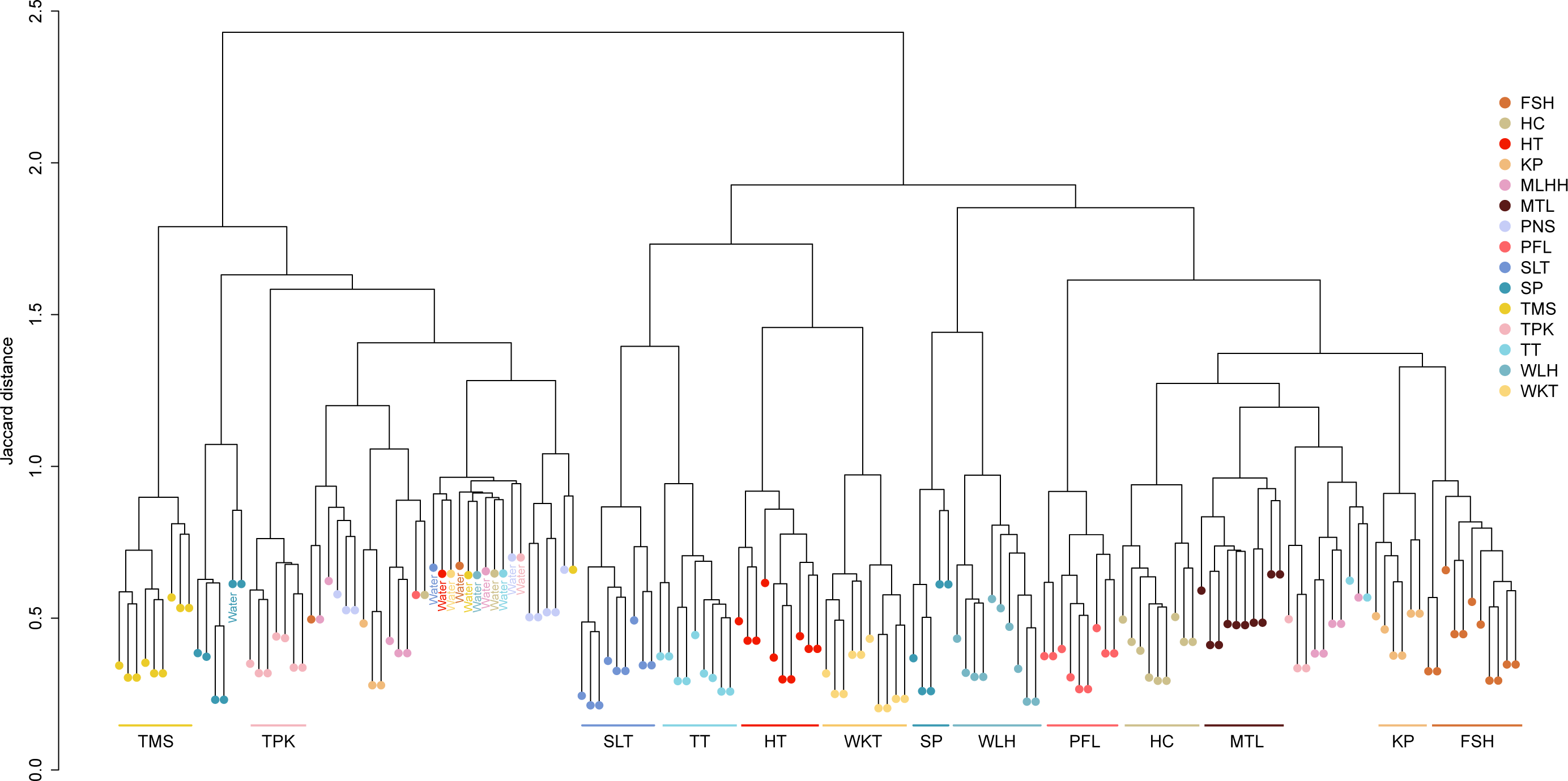
Hierarchical clustering of *P. hongkongensis* skin microbiomes from 15 sampling sites based on Jaccard distance. Clusters with at least five samples from the same site are indicated by the site name. Refer to Fig. 1 for the abbreviations of sampling sites.

